# Signaling through polymerization and degradation: Analysis and simulations of T cell activation mediated by Bcl10

**DOI:** 10.1101/2020.05.28.120907

**Authors:** Leonard Campanello, Maria K. Traver, Hari Shroff, Brian C. Schaefer, Wolfgang Losert

## Abstract

The adaptive immune system serves as a potent and highly specific defense mechanism against pathogen infection. One component of this system, the effector T cell, facilitates pathogen clearance upon detection of specific antigens by the T cell receptor (TCR). A critical process in effector T cell activation is transmission of signals from the TCR to a key transcriptional regulator, NF-κB. The transmission of this signal involves a highly dynamic process in which helical filaments of Bcl10, a key protein constituent of the TCR signaling cascade, undergo competing processes of polymeric assembly and macroautophagy-dependent degradation. Through computational analysis of three-dimensional super-resolution microscopy data, we quantitatively characterized TCR-stimulated Bcl10 filament assembly and length dynamics, demonstrating that filaments become shorter over time. Additionally, we developed an image-based bootstrap-like resampling method to quantitatively demonstrate preferred association between autophagosomes and Bcl10-filament ends and punctate-Bcl10 structures, implying that autophagosome-driven macroautophagy is directly responsible for Bcl10 filament shortening. We probe Bcl10 polymerization-depolymerization dynamics with a stochastic Monte-Carlo simulation of nucleation-limited filament assembly and degradation, and we show that high probabilities of filament nucleation in response to TCR engagement could provide the observed robust, homogeneous, and tunable response dynamic. Furthermore, the speed of autophagic degradation of filaments preferentially at filament ends provides effective regulatory control. Taken together, these data suggest that Bcl10 filament growth and degradation act as an excitable system that provides a digital response mechanism and the reliable timing critical for T cell activation and regulatory processes.

**Author Summary:** The immune system serves to protect organisms against pathogen-mediated disease. While a strong immune response is needed to eliminate pathogens in host organisms, immune responses that are too robust or too persistent can trigger autoimmune disorders, cancer, and a variety of additional serious human pathologies. Thus, a careful balance of activating and inhibitory mechanisms are necessary to prevent detrimental health outcomes of immune responses. For example, activated effector T cells marshal the immune response and direct killing of pathogen-infected cells; however, effector T cells that are chronically activated can damage and destroy healthy tissue. Here, we study an important internal activation pathway in effector T cells that involves the growth and counterbalancing degradation (via a process called macroautophagy) of filamentous cytoplasmic signaling structures. We utilize image analysis of 3-D super-resolution images and Monte Carlo simulations to study a key signal-transduction protein, Bcl10. We found that the speed of filament degradation has the greatest effect on the magnitude and duration of the response, implying that pharmaceutical interventions aimed at macroautophagy may have substantial impact on effector T cell function. Given that filamentous structures are utilized in numerous immune signaling pathways, our analysis methods could have broad applicability in the signal transduction field.

## Introduction

The vertebrate adaptive immune system consists of billions of T and B lymphocytes, which serve as a potent and highly specific host defense mechanism against pathogen infection. Once an antigen, an immune-activating component of a pathogen, has been recognized by the T cell receptor (TCR), an intracellular signaling cascade is initiated that leads to the clonal expansion of T cells responsive to this antigen. This anti-pathogen immune response, including the T cell component, will cause damage to healthy tissue if not temporally limited (1). Indeed, unregulated adaptive immune responses have been implicated in the development of certain cancers (1) and autoimmune diseases (2). Thus, adaptive immune responses include programmed shutdown mechanisms to maintain homeostasis and prevent damage to host tissues (2,3). In this study, we introduce novel image-analysis tools to robustly analyze a component signaling pathway of the adaptive immune response from a biophysical perspective, specifically focusing on the spatial organization of a key signaling structure that contributes to adaptive immunity.

The main cell types controlling adaptive immunity are the many classes of effector T cells, terminally differentiated T lymphocytes with previous antigen exposure, which can generate rapid anti-pathogen responses. Antigen engagement of the TCR in effector T cells initiates an internal signal transduction pathway that leads to the translocation of NF-κB from the cytoplasm to the nucleus. NF-κB is a heterodimeric protein complex that controls gene transcription in response to a diverse array of receptors (4) in both vertebrates and invertebrates (5). In effector T cells, the nuclear translocation of NF-κB results in the de novo or increased expression of a large number of genes involved in T cell proliferation and immune response mechanisms (5,6). Improper regulation of NF-κB signaling in T cells is mechanistically associated with primary immunodeficiency (7–9) and autoimmune disease (10,11). Thus, the regulation of the TCR-to-NF-κB pathway is critically important for proper immune function.

A key signaling protein in the TCR-to-NF-κB pathway is Bcl10 (12). Upon TCR stimulation, Bcl10 assembles with its signaling partners Carma1 and Malt1 to form the micron-scale ‘CBM’ complex (13–17). In effector T cells, the CBM complex forms the core of a filamentous assembly called the POLKADOTS signalosome, which serves as the cytoplasmic site of the terminal steps of the NF-κB-activation cascade (13,14,18–21). Contemporaneously with assembly into microns-long filaments, Bcl10 is also degraded (13,22–26), and we have previously shown that proteolysis of Bcl10 occurs within the POLKADOTS signalosome via TCR-dependent selective autophagy (18). In this degradative process, autophagosomes associate with POLKADOTS filaments resulting in selective destruction of Bcl10, thus terminating signals to NF-κB. Data suggest that the balance between assembly of filamentous Bcl10 in POLKADOTS and proteolytic degradation of Bcl10 via selective autophagy determines the extent of NF-κB activation (13,18).

Notably, many outputs of TCR signaling exhibit nonlinear response profiles, including an all-or-nothing (i.e., digital) responses. While not every T cell response has digital characteristics (27), studies have demonstrated that specific TCR-triggered events are digital, including activation of the extracellular regulated kinase-mitogen activated protein kinase (ERK-MAPK) signaling cascade (28–30), signaling via the Protein kinase D2-protein kinase C (PKD2-PKC) cascade (31), the release of cytokines (32), and the activation of cytolytic capacity (33). Additionally, we have previously shown that TCR activation of NF-κB is digital in nature, with formation of and signal transmission by the POLKADOTS signalosome occurring in an all-or-nothing manner (34). Mechanistically, how nucleation, growth and eventual autophagic degradation of POLKADOTS filaments are connected to the digital nature of the NF-κB cascade is unclear. In this manuscript, we link the nonlinear character of the signaling cascade to the spatial organization, assembly, and degradation of filamentous Bcl10. We propose that highly nonlinear self-assembly and degradation processes serve as an excitable system enabling a rapid, and at the same time, self-limiting response.

Several key details of this polymerization process are known: recent cell-free cryo-electron-microscopy studies revealed that Bcl10 polymerizes into filaments with a geometrically confined, helical arrangement with other POLKADOTS components (14,20,21). Although the core of the POLKADOTS filament is composed primarily of Bcl10, polymerization of Bcl10 is nucleated by short oligomers of Carma1and the Carma1-driven nucleation of Bcl10 can act as an amplifier of activation signals from the T cell receptor to NF-κB (14). In contrast to the above parameters driving filament growth and signal propagation, autophagic degradation of Bcl10 opposes this amplification process; however, the geometric and kinetic features that limit and ultimately reverse filament growth in living T cells are unclear. These opposing effects on filament growth and degradation are not well understood and cannot be assessed via traditional biochemical approaches and signaling pathway models, which excludes spatial organization.

In this study, we quantify the spatial arrangement of Bcl10, its colocalization with autophagosomes, and the randomness of their relative spatial distributions. Our analysis introduces a novel computational workflow that combines bootstrap-like resampling methods adapted for image features and medial-axis-thinning skeletonization to analyze the structure and organization of Bcl10 filaments in relation to autophagosomes in a robust manner. The methods we present are broadly applicable to studies of biologically encoded spatial colocalizations and self-assembly, such as toll-like receptor (TLR)-stimulated assembly of the myddosome, RIG-I-like receptor (RLR) triggering of mitochondrial antiviral-signaling protein (MAVS) oligomerization, and activation of the various sensor proteins that promote assembly of micron-scale inflammasomes (35). To gain further mechanistic insights into the assembly and degradation dynamics of POLKADOTS filaments, we complemented analysis of super-resolution images with stochastic Monte Carlo simulations of POLKADOTS dynamics that include the nucleation, growth, and degradation of Bcl10. In sum, we show that spatial assembly and degradation of protein complexes in the TCR-to-NF-κB-activation cascade are key design elements to ensure a rapid response yet prevent ongoing immune activation. These elements may also contribute to the digital nature of this crucial signaling pathway.

## Data Availability Statement

The raw data, processed data, analyzed data are available via Google Drive.

## Results

### Measuring Bcl10 Filament Lengths

Utilizing a long-term murine effector T cell clone, D10.G4.1 (henceforth referred to as D10), we engineered a cell line stably expressing GFP-tagged Bcl10, which we used to image Bcl10 filaments in cells fixed at 20- and 40-min post-TCR activation using a super-resolution instant structured illumination microscope (iSIM) (36) (**Fig 1A**). The earliest evidence of formation of POLKADOTS structures is at 10 min post-TCR activation, and these polymers are maximally evident by 20 min post-activation, at which point initial activation with autophagosomes is also apparent. Association with autophagosomes and degradation of Bcl10 persists through 2 hr post-stimulation, by which time Bcl10-containing polymers can no longer be found in any cells (and these structures are rare even by 60 min post-activation) (13,15). Thus, 40 min post-activation is a reasonable time point to assess intermediate consequences of autophagosome association with Bcl10 filaments.

**Fig 1.**
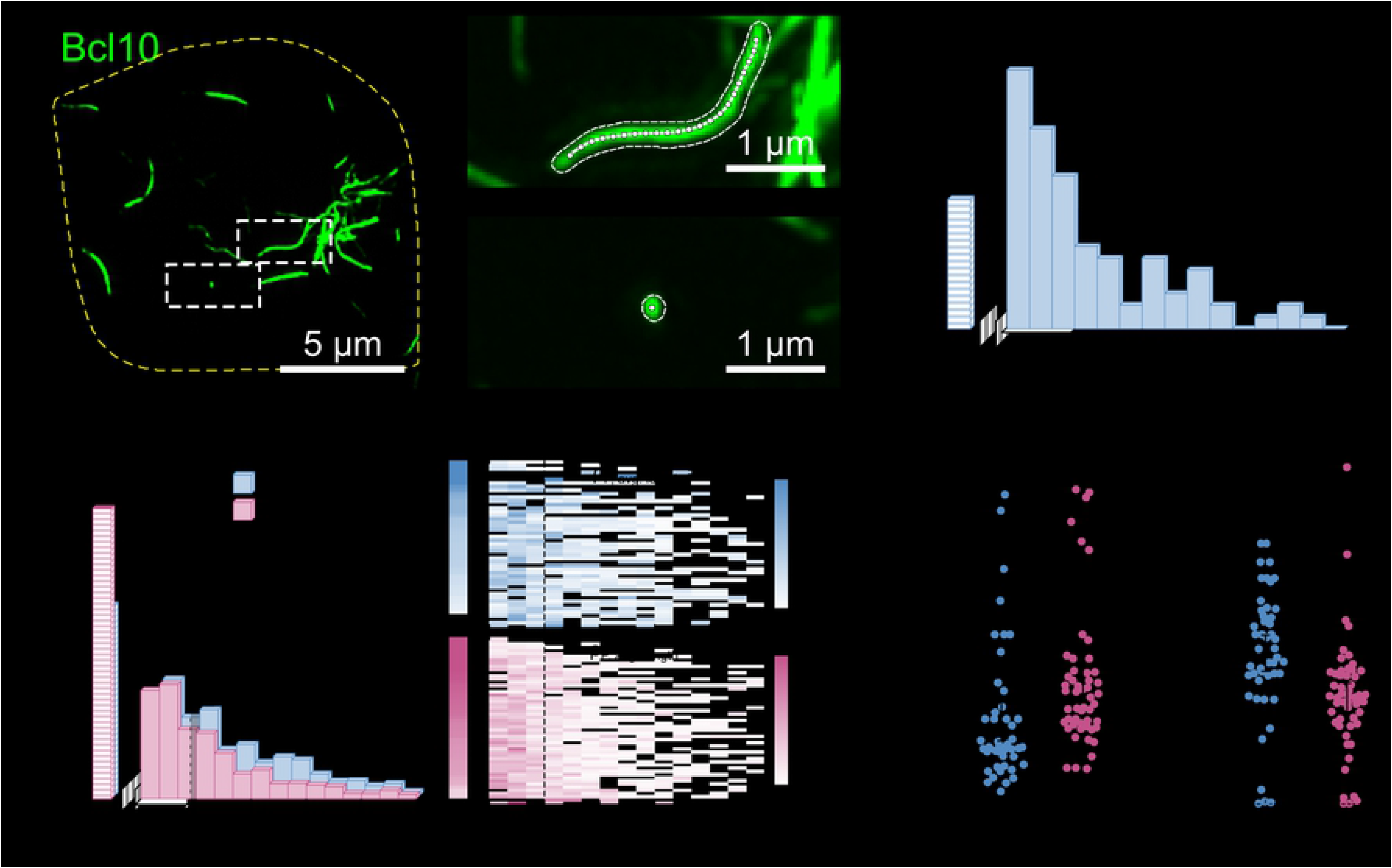
Bcl10 filaments shorten between 20 minutes and 40 minutes post-T cell receptor activation. (A) Representative image (maximum intensity projection) of an activated T cell 20 minutes after TCR stimulation. The outlined region of interest contains two complete Bcl10 filaments. (B) Representative Bcl10 filaments within the activated T cell. Filament outlines and skeletons were calculated using a semiautomatic segmentation method and medial axis thinning. The upper image exemplifies the filamentous form of Bcl10 and the lower image is representative of punctate Bcl10. (C) Representative distribution of Bcl10 lengths from all Bcl10 filaments from A. Punctate Bcl10 structures were separately counted and binned as “P”. (D) Cumulative length distributions of all Bcl10 filaments in all T cells at 20 minutes (N = 47 cells from 2 independent experiments) after TCR stimulation and 40 minutes (N = 53 from 2 independent experiments). Rows of the heat map are the length distributions for cells, sorted by the percentage of Bcl10 structures in each cell that are punctate. The correlation between the number of punctate structures and average filament length is −0.657 at 20 minutes post-activation, and −0.695 at 40 minutes post-activation. (E) Scatter plot of the proportion of Bcl10 structures which are punctate at 20 minutes and 40 minutes after TCR stimulation. (F) Scatter plot of the proportion of Bcl10 filaments longer than 1 μm at 20 minutes and 40 minutes after TCR stimulation. Correlations are the Pearson product-moment correlation. Error bars represent the 95% confidence interval of the mean. *p*-values were calculated using a two-sample *t*-test. *p*-values less than 0.05 were considered significant.

To analyze these data, we developed a semiautomatic segmentation and skeletonization algorithm based on medial-axis thinning, then utilized the resultant skeletons to obtain various structural measurements (**Methods** and **SI Movie S1**). Bcl10-rich regions with skeleton-lengths <150 nm (the approximate spatial resolution of the iSIM) were designated as punctate structures, labeled by “P” in these data, and analyzed separately from micron-scale filaments (**Fig 1B**), which ranged in length from 0-5 μm (**Fig 1C**). Between 20 minutes and 40 minutes post-activation, we observed an increase in the relative number of Bcl10 puncta, along with a corresponding decrease in the number of long Bcl10 filaments (**Fig 1D**). Furthermore, at both timepoints, a moderate negative correlation (ρ < −0.6) existed between the overall number of Bcl10 puncta and the distribution of Bcl10 filament length; thus, cells with fewer puncta were more likely to have long filaments, and cells with more puncta were more likely to have shorter filaments (**Fig 1D**). This correlation existed and was statistically significant at the single cell level; cells at 40 min post-stimulation were more likely to have both a larger relative proportion of punctate structures (**Fig 1E**, **SI Table S2**, p = 1.0 × 10^−3^), and a decrease in the relative number of Bcl10 filaments longer than 1 μm (**Fig 1F**, **SI Table S2**, p = 3.9 × 10^−4^). Taken together, these results are consistent with the interpretation that Bcl10 filaments decrease in length between 20 min and 40 min post-activation, and that the proportion of punctate structures increases. This increase in the number of punctate structures and decrease in the number of long filaments could indicate (i) ongoing nucleation of new filaments, (ii) disassembly or end-directed degradation of existing filaments, (iii) scission of existing filaments, which would create two “daughter” filaments with each scission event, or some combination of the above processes.

### Examining Autophagosome-Bcl10-Filament Interactions

Our previous work has demonstrated that Bcl10 within the filamentous POLKADOTS signalosome is targeted for degradation by macroautophagy (henceforth referred to as autophagy) (18), an intracellular degradative process involving the envelopment of cargo by double-membraned vesicles called autophagosomes. Autophagic degradation of Bcl10 leads to a dampening of NF-κB activation and NF-κB-dependent T cell responses (18). To examine whether autophagy is responsible for the observed decrease in Bcl10 filament length, we expressed an RFP-tagged form of the autophagosome membrane protein LC3 in our Bcl10-GFP-expressing D10 cell line, then imaged static interactions between Bcl10 filaments and LC3-labeled autophagosomes at 20- and 40-min post-TCR activation (**Fig 2A**). Consistent with our previous data (18), our experiments indicated that filamentous Bcl10 and LC3-positive autophagosomes existed simultaneously in activated T cells. Autophagosome contacts with Bcl10 filaments were frequently observed; thus, independent semiautomatic segmentation of Bcl10 filaments and autophagosomes was used to determine the number and location of contacts (**Fig 2B**). Interestingly, we observed that activated T cells have significantly fewer Bcl10-autophagosome contacts at 20 min post-TCR stimulation than at 40 min post-stimulation (**Fig 2C**, **SI Table S3**, p = 1.45 × 10^−4^); however, this fact alone is not sufficient to conclude that contact formation is preferred. To assess whether autophagosome contacts with Bcl10 filaments form preferentially or randomly, we developed an image-based-resampling method analogous to statistical bootstrapping (**Methods**). We randomly rearranged the segmented autophagosomes throughout the cell cytosol and recalculated the resulting filament-autophagosome contacts (**Fig 2D**). The cell-averaged number of colocalizations in these rearrangements was significantly fewer than the number in the experimental observations (**Fig 2E**). As an additional test, we conducted 100 rearrangements of each individual cell and constructed a confidence interval for the number of randomized contacts. The actual number of autophagosome contacts in 45 of 47 cells at 20 minutes post-activation, and 52 of 53 cells at 40 minutes post-activation, was greater than the 95% confidence interval for random LC3 contacts (**Fig 2F**). Thus, we concluded that contact between Bcl10 and autophagosomes was non-random, and there were indeed more Bcl10-autophagosome interactions at 40 minutes post-activation.

**Fig 2.**
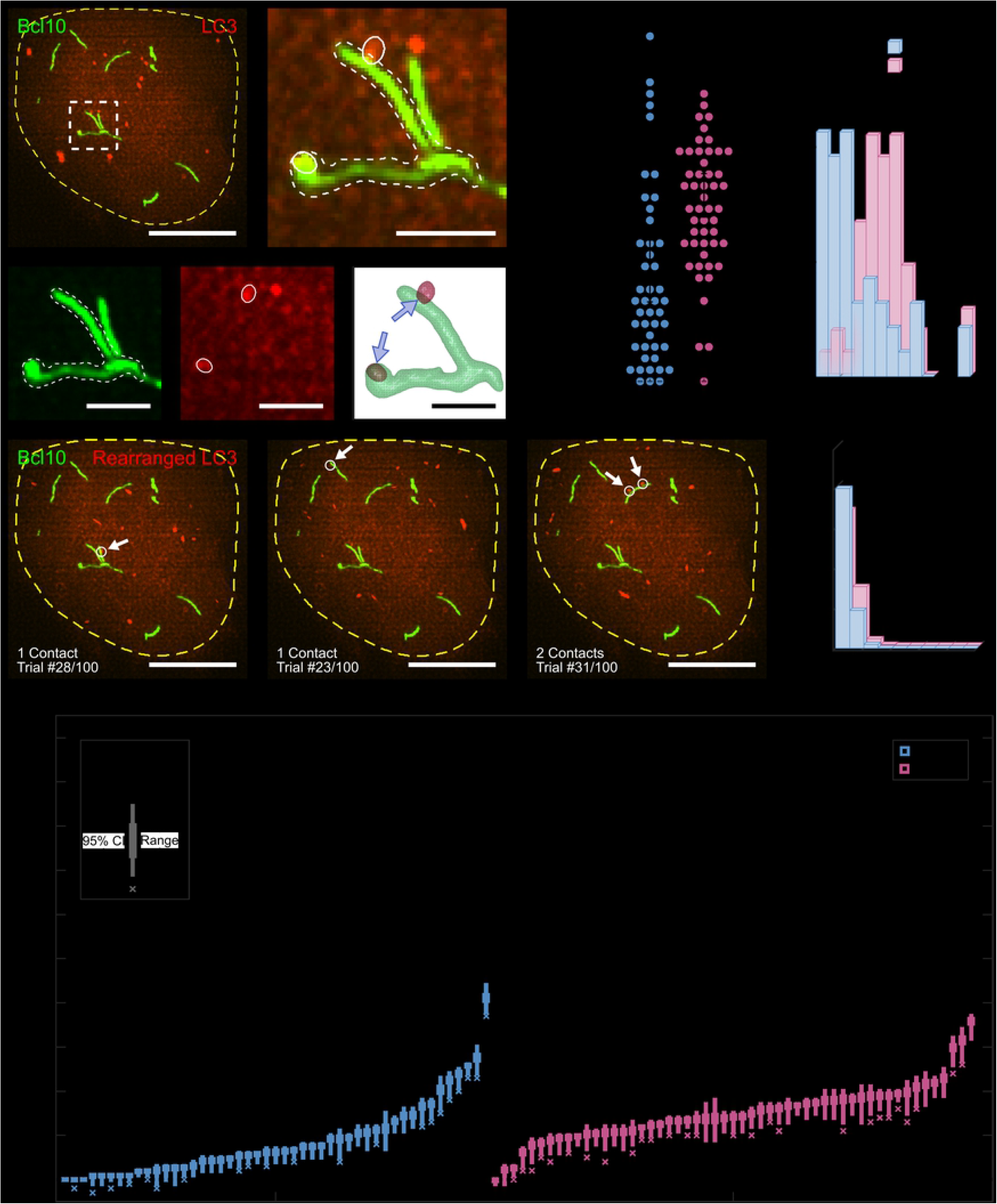
Bcl10 filaments are preferentially in contact with autophagosomes. (A) Representative image of an activated T cell 20 minutes after TCR stimulation. Green Bcl10 filaments and red LC3 vesicles (autophagosomes) are within the yellow cell boundary. The outlined region of interest contains a Bcl10 structure with two colocalized autophagosomes. (B) Representative Bcl10 filament-autophagosome contacts. The semiautomatic segmentation and skeletonization analyses identify the number and location of colocalizations which are highlighted by blue arrows. (C) Cumulative number of Bcl10 filament-autophagosome contacts in all T cells (20 min: N = 47 cells from 2 independent experiments; 40 min: N = 53 cells from 2 independent experiments). There is a statistically significant difference in the number of Bcl10-autophagosome contacts at 20-min- and 40-min-post-stimulation (p = 1.4 × 10 ^‒ 4^). (D) Three representative trials of the random-spatial rearrangement of autophagosomes in the representative cell from A. Contacts that formed after rearrangements are highlighted with arrows. (E) The cumulative distribution of the number of Bcl10 filament-autophagosome contacts in the random rearrangement trials. (F) Difference between the true number of Bcl10 filament-autophagosome contacts and the number of contacts found in each of the 100 trials for each cell. Positive values indicate that there are more true contacts than what is found in the trials. There is statistically significant difference in the number of contacts between the true and rearranged data in 45 of 47 T cells imaged 20 min post-activation, and 52 of 53 T cells imaged 40 min post-activation. Error bars are the 95% confidence interval of the mean and were calculated independently for each cell. *p*-values were calculated using two-tailed *t*-tests. *p*-values less than 0.05 were considered significant.

Our results thus far indicate that at 40 minutes post-activation, Bcl10 filaments are both shorter and have increased contact with autophagosomes, consistent with the interpretation that autophagosomes are degrading the filaments over time. However, a mechanistic problem arises from this data: autophagy occurs when double-membraned vesicles completely surround and envelop intracellular cargo, yet Bcl10 structures are larger than autophagosomes–the majority of autophagosome structures are smaller than (0.2 μm)^3^ in volume at both 20 min and 40 min post-activation (**SI Fig S4**). Thus, the traditional biophysical understanding of mechanisms of autophagy may not be viable as an explanation for Bcl10-filament degradation. To examine the mechanism whereby autophagosomes degrade Bcl10 filaments, we examined the location of autophagosome colocalizations along each filament. Biochemical and structural studies have demonstrated that Bcl10 filaments have two distinct end points and no junctions along the main body (14); however, due to the resolution limits of this imaging method, some filaments appear joined.

Thus, employing the same super-resolution dataset used for **Fig 2**, we extracted Bcl10 filament skeletons and fragmented them to eliminate high-degree junctions such that the resulting skeleton configurations minimized the total bending energy (**Fig 3A**, **Methods**). After processing the skeletons, each Bcl10 filament was segmented based on the distance from the nearest skeleton endpoint (**Fig 3B**). Finally, we examined each colocalization event between an autophagosome and a Bcl10 structure; if the Bcl10 structure was punctate, the event was designated “P”, and if the Bcl10 structure was filamentous, the event was characterized by the shortest distance between the autophagosome and the skeleton endpoint (**Fig 3C**). Excluding puncta, the distance-from-end distributions at 20 min and 40 min were not statistically distinguishable (**Fig 3C**, **SI Table S5**, p = 0.085), and we found a marked preference for autophagosomes to localize near filament ends at both time points. Interestingly, a greater proportion of autophagosomes were in contact with Bcl10 puncta (as opposed to filaments) at 40 min post-activation than at 20 min post-activation (**Fig 3C**, **SI Table S5**, p = 2.03 × 10^−9^). Though the proportion of punctate Bcl10 approximately doubled between 20 and 40 min post-activation (**Fig 1E**), the proportion of autophagosomes that localized to puncta approximately quadrupled (**Fig 3C**), indicating a distinct preference for autophagosomes to localize with punctate Bcl10 at the later time point.

**Fig 3.**
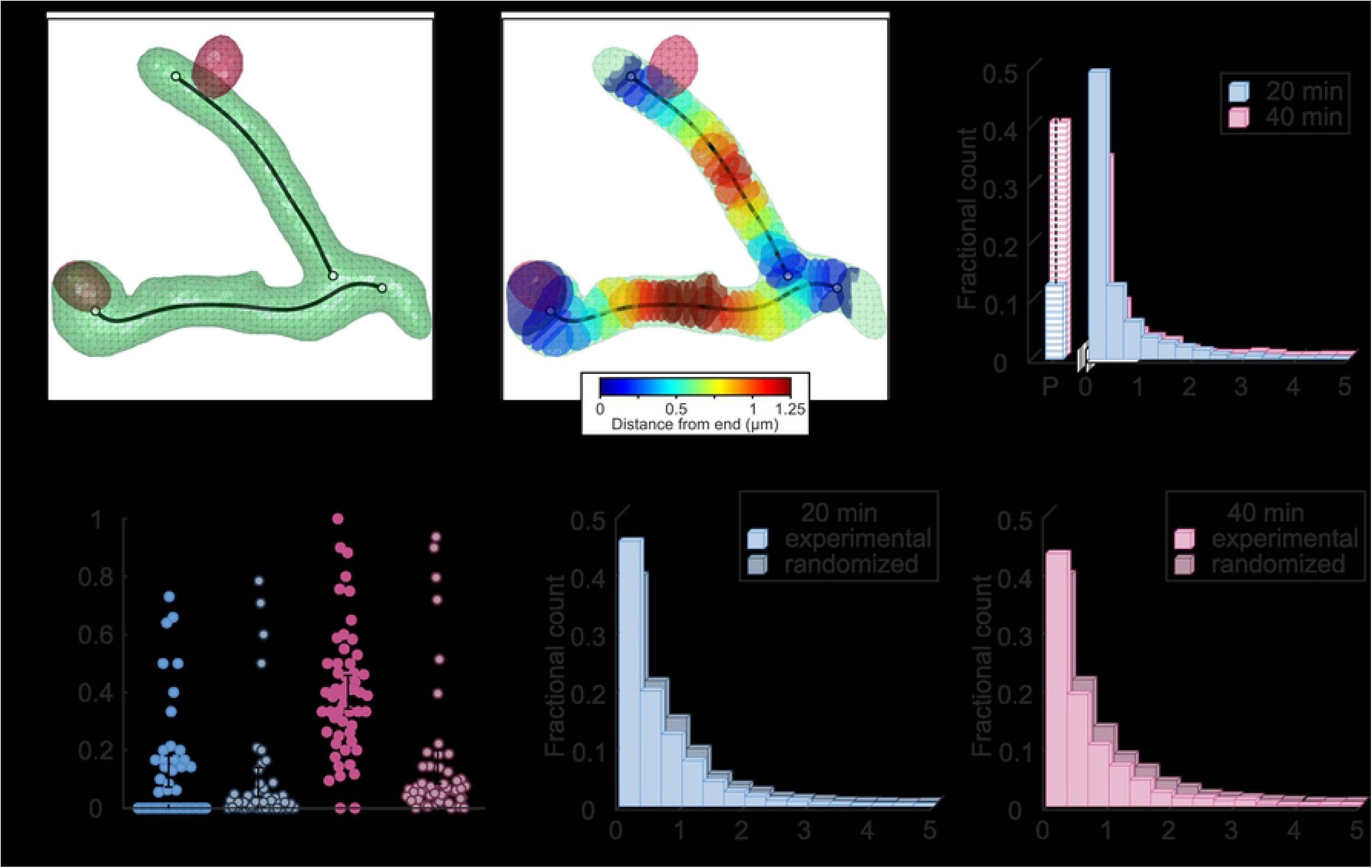
Autophagosomes preferentially localize to Bcl10-filament ends and, at later timepoints, to Bcl10 puncta. (A) Representative segmented Bcl10 filaments (green) and autophagosomes (red) after post-processing overlaid with the pruned medial-axis skeleton of Bcl10 (black). Filament skeletons were fragmented to minimize the bending energy at junctions. (B) Using the skeleton of the representative Bcl10 filament, the filament volume was segmented into sections based on the nearest distance from skeleton ends. (C) Distribution of autophagosome-Bcl10 contacts by distance from endpoint (filamentous Bcl10) or punctate Bcl10 (“P”). (D) Scatter-bar plot of the proportion of Bcl10 puncta per cell in contact with an autophagosome in experimental and randomized data. (E) Distribution of autophagosome-filamentous Bcl10 colocalizations by distance from endpoint in experimental and randomized data. *p*-values in C and E were calculated using two-sample KS tests and *p*-values in D were calculated using paired two-sample *t*-tests. *p*-values less than 0.05 were considered significant.

Our observations thus far indicated that autophagosomes were more likely to localize to the ends of filaments and, by 40 minutes post-activation, to Bcl10 puncta. To assess whether these localizations were truly preferred, or whether they were in fact random, we again utilized the randomized images from the image-based resampling shown in **Fig 2D**. First, we compared the number of colocalizations of Bcl10 puncta with autophagosomes in the true versus the randomized images (**Fig 3D**). At 20 minutes post-activation, the true and random distributions of contacts were indistinguishable (**Fig 3D**, **SI Table S5**, p = 0.079), indicating that autophagosome-puncta contacts were likely the result of random co-localization. However, at 40 minutes post-activation, the number of true puncta contacts was greater than the number of such contacts in the randomized images, and this enrichment was statistically significant (**Fig 3D**, **SI Table S5**, p = 2.15 × 10^−16^), indicating that punctate Bcl10 structures were preferentially in contact with autophagosomes at this time point. Next, we compared the distance-from-end distributions of autophagosome-filament contacts for the experimental versus randomized data at both time points (**Fig 3E**). The randomized distributions at both time points demonstrated fewer end-localizations and increased numbers of contacts along the body of the filament in comparison to quantification of localization data from true images (**Fig 3E**, **SI Table S5**, p = 1.10 × 10^−11^ and 2.01 × 10^−16^ at 20 min and 40 min, respectively). These results indicate that the preference for autophagosomes to localize at or near filament ends was non-random.

Taken together, the results from **Fig 2** and **Fig 3** indicate that following TCR engagement and activation, autophagosomes formed attachments with Bcl10 filaments, the number of attachments increased over time, and these attachments occurred near the ends of Bcl10 filaments. Furthermore, by 40 minutes post-activation, autophagosomes also formed attachments with Bcl10 punctate structures in a biologically targeted (i.e., non-random) manner. Together, these results are consistent with the hypothesis that autophagosomes are responsible for the progressive shortening of Bcl10 filaments over time, and that this process is directed by a biologically encoded targeting process. Further, the accumulation of punctate Bcl10 structures at 40 minutes post-activation may be the result of the degradation of longer Bcl10 filaments. Complete degradation of Bcl10 may follow a non-linear kinetic, with the final remnant, visualized as puncta, undergoing slower degradation than filamentous regions of these structures.

### Modeling Bcl10 Filament Dynamics

Our previously published observations demonstrate that macroautophagy contributes to Bcl10 filament degradation (18), and the above analysis of imaging data support and extend these conclusions. This degradation occurs simultaneously with Bcl10 polymerization, and the interplay between these processes serves to control T cell activation states. However, our understanding of the dynamics of filament polymerization and degradation, and the interaction between these processes, remains limited.

To further explore the interplay between simultaneous Bcl10 polymerization and degradation, and its implications for control of T cell activation states, we designed a stochastic Monte Carlo simulation to model Bcl10 polymerization (**Fig 4** and **Methods**). Previous studies have indicated that growth of Bcl10 filaments is nucleated by short cytosolic oligomers of Carma1 (14,20), and that this nucleation is the rate-limiting step for filament growth (14); thus, we used nucleation-limited filament assembly as the basis for our simulation. Nucleation-limited assembly is similar to a Bernoulli process whose probability parameter increases when the length of the growing filament reaches a critical length: the size of the nucleation barrier. Under these conditions, theoretical predictions of filament growth rate indicate a dramatic transition between slow, initial growth, to fast, steady-state growth (**SI Text S6**). Further, filament growth is mediated by a kinetic process whereby monomers collide with or are recruited to the site of the growing filament. Our simulation began with the generation of “cells,” which were each given a random size that was used to generate the initial conditions, including numbers of monomeric nucleation-site components (mimicking monomeric Carma1), Bcl10, autophagosomes, and the size of the immunological synapse (i.e., the portion of the cell membrane with the potential to activate the Bcl10-nucleation sites). As this cell evolved in time over each iteration of the simulation, the growth of Bcl10 filaments was governed by three separate independent and tunable parameters (visually represented in **Fig 4A**): the activation probability of the nucleation sites (p_activate_), the growth probability of the initial layer of Bcl10 monomers before the nucleation barrier has been overcome (p_attach_), and the steady-state growth probability after the filament overcomes the nucleation barrier (p_grow_).

**Fig 4.**
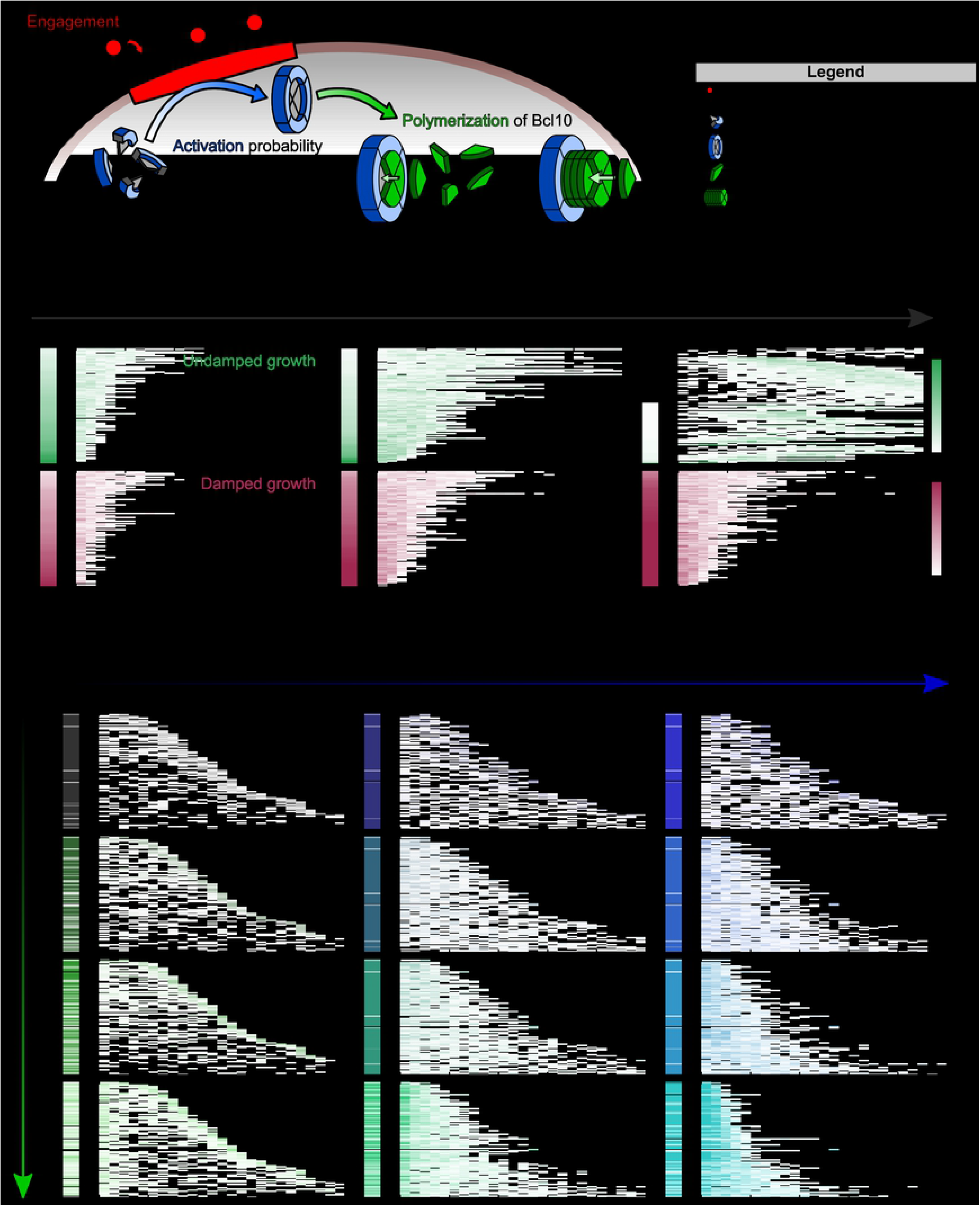
Simulations of Bcl10 filament growth reveal properties of real Bcl10 nucleation rates and concentrations, but fail to match observed values for ρ. (A) Simulated Bcl10-filament growth is controlled by three parameters: the nucleation-site activation probability (p_activate_), the initial layer attachment probability (p_attach_), and the steady state attachment probability (p_grow_). (B) Snapshots of representative simulations of Bcl10 filament growth in undamped (green, top) and damped (red, bottom) environments at 10%, 20% and 100% of n_max_. Each horizontal line in the kymographs represent a simulated cell with its own unique set of initial conditions, including number of nucleation-site components and initial Bcl10 concentration, and each simulation was terminated after n_max_ iterations, which is the number of iterations for the number of free Bcl10 monomers to reach zero. (C) Changing simulation parameters exposes different behaviors in a heterogeneous population of T cells. Here, the nucleation-site-activation and initial-layer-attachment probabilities are simultaneously modulated in the same heterogenous population of 100 T cells with varying numbers of nucleation sites and initial Bcl10 concentrations. From left to right, the activation rate of nucleation sites is increased. From top to bottom, the Bcl10 collision rate is increased. All simulations are terminated when the concentration of Bcl10 monomers reaches zero.

First, we sought to determine whether the value of each parameter should remain constant throughout each iteration of the simulation, or whether the results of each iteration would affect parameter values. For instance, if the concentration of free Bcl10 monomers decreases over time (i.e., if resynthesis of Bcl10 monomers is outpaced by their degradation), the probability of monomer collisions with the growing filament end decreases over time, thus decreasing p_grow_, a process known as damping. To determine whether Bcl10 filament growth is damped, we tested 100 simulated T cells, each with a different “size” and thus initialized with a unique number of nucleation sites and Bcl10 monomers, representing the evolution of a heterogeneous population of T cells with varying initial conditions (**Fig 4B**). When our simulation ran in an undamped environment, growth continued perpetually and, by the end of the simulation, punctate structures all grew into filaments and all Carma1-mimickingnucleation sites were filled. Running the same simulation in a damped environment, with a finite supply of Bcl10 monomers, led to a heterogeneous distribution of Bcl10 puncta and filaments of varying lengths. The results of the damped growth simulation were more representative of observed Bcl10 filament lengths (**Fig 1D**), leading us to conclude that Bcl10 filament growth is limited by Bcl10 monomer supply, a conclusion supported by biochemical analyses of Bcl10 levels over time in activated T cells (18). Thus, in all subsequent simulations, we allowed the decreasing availability of monomeric Bcl10 to cause an effective stochastic reduction in the Bcl10-nucleation probabilities, p_attach_ and p_grow_, since fewer monomeric Bcl10 will lead to a reduced number of attempted attachments.

Next, we examined whether changes in the value of p_grow_ would provide useful information on the response dynamics. Previous work by David et al. (14) indicated that nucleation (p_activate_) and/or the initial attachment of Bcl10 filaments (p_attach_) is the rate-limiting step in filament growth. Thus, varying the value of p_grow_ would likely provide little in the way of relevant biological information. Further, increasing the growth rate would more quickly deplete the monomer concentration, thus ending the simulation after fewer iterations and limiting the sampling of the activation and attachment probabilities. Thus, we chose to assign p_grow_ a fixed value that allowed the simulations to run for a reasonable number of iterations (here, we chose p = 0.4 as described in the **Methods**), and we only modulated p_activate_ and p_attach_.

Finally, we examined the effect of various values for p_activate_ and p_attach_ on filament lengths resulting from the simulation (**Fig 4C**). The observed filament lengths in our cell images (**Fig 1D**) demonstrate moderately negative correlations between the relative numbers of puncta and length of Bcl10 filaments, ρ < −0.6; thus, we examined which combination of parameters applied to the same heterogenous population of 100 simulated T cells could generate an equivalent correlation. Each simulation was repeated 10 times, and each trial was terminated when the number of free monomers of Bcl10 reached zero; the filament-length distributions we report represent the cumulative distribution from each of the 10 trials. There was complex interplay between p_activate_ and p_attach_: when p_activate_ was low, there were few simulated filaments and simulated filament length distributions appear to be unchanged with increasing p_attach_ (**Fig 4C – left**). On the other hand, when p_activate_ was high, there were many activated nucleation sites in each simulated cell, which resulted in heterogeneity in the preferred filament length: increased p_attach_ resulted in decreased average-filament length (**Fig 4C – right**). Interestingly, intermediate values of p_activate_ in the simulations result in non-linear behavior (**Fig 4C – center**): the preferred filament length appeared to be independent of p_attach_ for small p_attach_, similar to **Fig 4C – left**, but decreased for large p_attach_, similar to **Fig 4C – right**. Careful tuning of simulations parameters resulted in correlations between ρ = [−0.5, −0.4] in subsections of the simulations-parameter space where p_activate_ was 0.5, and p_attach_ was 0.075, but the simulation was unable to reach observed levels of ρ < −0.6. This failure to match the model to experimental observations points to a need to include Bcl10 degradation in the simulations as an important contributor to the process.

To understand the effect of degradation on Bcl10 filament size, we added p_degrade_, which represents the probability in the simulations that an autophagosome will attach to a filament and cause degradation (**Fig 5A**). As in **Fig 4**, we simulated 100 cells with random initial conditions and repeated each simulation 10 times. A comparison of twelve representative parameter sets is instructive to determine the effects of each individual parameter, as well as their interplay, on the concentration of filamentous Bcl10 (**Fig 5B**). For this set, we chose a low vs a high value for both p_attach_ (**Fig 5B, top vs bottom**) and p_activate_ (**Fig 5B, left vs right**); for each of these four parameter subspaces, we then compared a low, medium, and high value for p_degrade_, which was varied over the same range as p_attach_. As expected, increasing values of p_degrade_ in each parameter space led to a smaller concentration of filamentous Bcl10. Interestingly, an increase in the rate of nucleation-site generation (represented by a greater p_activate_) led to a faster decline in filamentous Bcl10 (**Fig 5B, left vs right**). This result makes intuitive sense, as a greater number of activation sites would lead to a greater number of filaments, and thus a greater number of sites where degradation can occur. In contrast, a lower activation barrier (represented by a greater p_attach_) had a much smaller effect on the rate of decline in filamentous Bcl10, promoting only a slight increase in filament stability. When combined, a lower activation barrier and a greater activation of nucleation sites created a dynamic in which slight variations in degradation had a large effect on the concentration of filamentous Bcl10.

**Fig 5.**
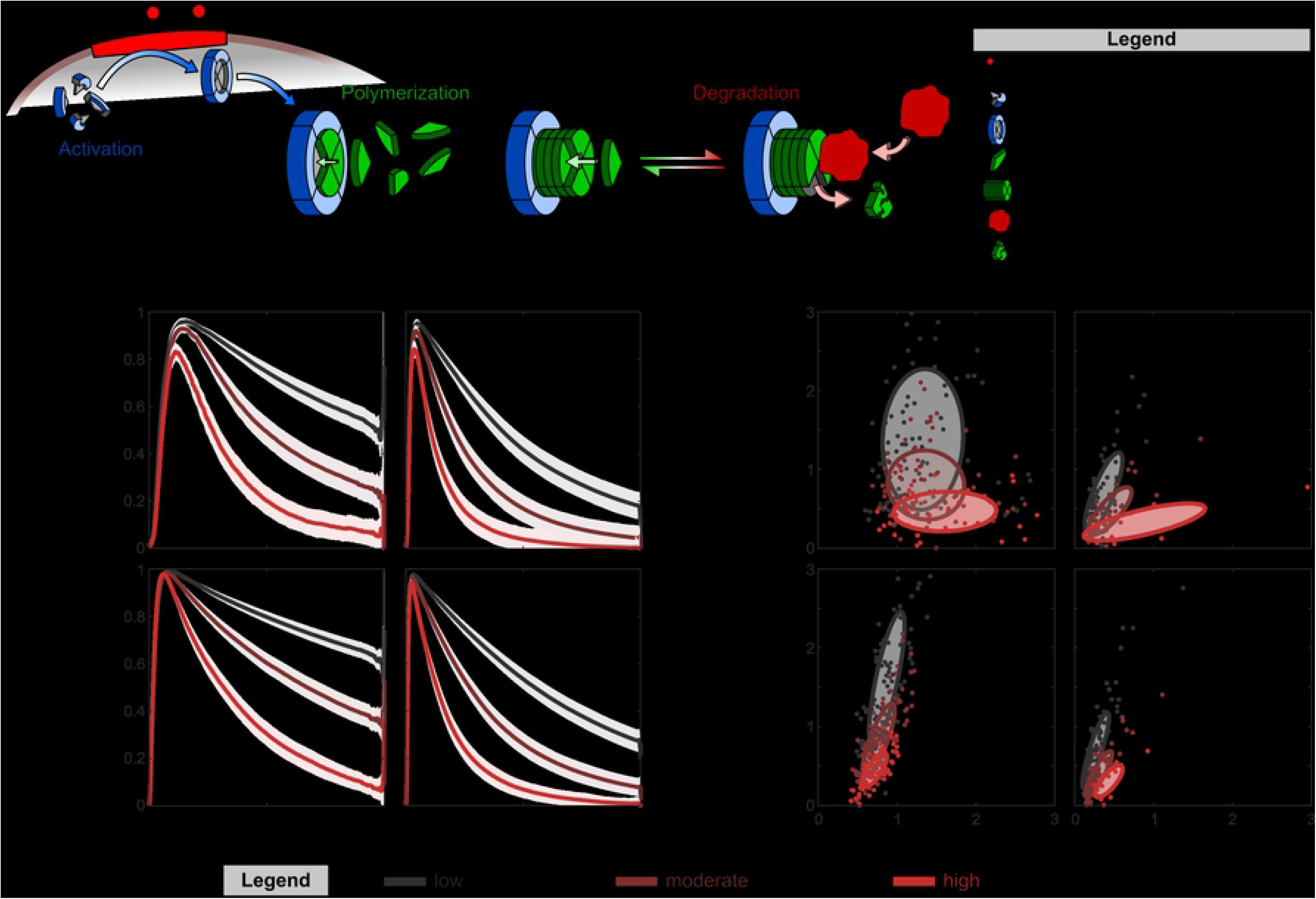
Simulations of Bcl10-filament growth and decay demonstrate that changes in the response dynamic due to increased degradation by autophagy were stabilized by increased p_attach_ but emphasized by increased p_activate_. (A) Simulated Bcl10-filament degradation was simulated in conjunction with filament growth. Degradation was modulated by p_degrade_, the probability of degradation by an autophagosome. (B) The average normalized concentration of filamentous Bcl10 as simulations evolved. Concentration is relative to the maximum-average concentration of all simulations with the same parameters. Each line represents the average of 100 cells, repeated 10 times each, and shaded bars are the 95% CI of the mean. Simulations were repeated with low and high values for p_activate_ and p_attach_. (C) The number of iterations needed to reach the maximum relative concentration vs. the number of iterations to reach half of the maximum concentration due to degradation. Each entry represents the average n_max_ and n_half_ for the 10 trials of each of the 100 cells. Low p_degrade_ is 0.05, moderate p_degrade_ is 0.10, and high p_degrade_ is 0.20. Ellipses represent one standard error of the mean.

We further explored how each of the three parameters in our simulation affected the variability of the response across the heterogeneous initial population of cells (**Fig 5C**). For each of the 100 cells, we calculated the average number of iterations for the filament-length distribution to reach its maximum average size, n_max_, and compared this to the average number of iterations required for the average size to be half of the max, n_half_ (**Fig 5C**), for each of the twelve parameter configurations. Increasing the value of p_attach_, p_activate_, or p_degrade_ independently of the others led to decreased variability in outcomes, with a lower nucleation barrier (an increase in p_attach_) having the greatest effect on variability in the response to an activation signal (i.e., heterogeneity in the time to peak response), and an increase in p_activate_ emphasizing changes in p_degrade_. The introduction of degradation into the simulations increased the negative correlations between the number of puncta and average filament length, ρ. Each of the simulations reached values of ρ ≈ −0.5 during system evolution, close to the experimental value of ρ < −0.6.

In summary, we have developed a refined model of a critical T cell receptor signaling process that quantitatively captured the growth and decay dynamics of Bcl10 filaments and their heterogeneous length distribution. We determined that a high activation potential of nucleation sites can, when combined with a low Bcl10-nucleation barrier, provide the most predictable and homogenous response to T cell activation. Furthermore, we found that the rate of degradation of Bcl10 filaments had the largest effect on the percentage of Bcl10 in filamentous form when nucleation sites have a high activation potential and Bcl10 has a low nucleation barrier. These conditions ultimately allow the response dynamic to be maximally tunable via changes in the degradation rate. With regards to the biological governance of this signal transmission system, these data offer a mechanistic explanation for why Bcl10 polymerization is opposed by contemporaneous autophagic degradation of Bcl10 polymers.

#### Supporting Information Legends

**SI Movie S1: Semiautomatic segmentation, skeletonization, and pruning of Bcl10 filaments.**

Representative segmented Bcl10 filament superimposed medial-axis skeletons. (Left) Raw medial-axis skeleton. The skeleton contains several small spurs which occur because of voxel noise on the surface of the filament. (Middle) Medial-axis skeleton after moderate pruning. Some spurs are removed. (Right) Medial-axis skeleton after full pruning. All spurs are removed and only the main-body skeleton remains.

**SI Table S2. Statistics for Fig 1.**

**SI Table S3. Statistics for Fig 2C.**

**SI Fig S4. Shape statistics of autophagosomes 20 min and 40 min post-activation.**

(A) Distribution of autophagosome 3-D volumes. (B) Distribution of autophagosome 3-D surface areas. (C) Distribution of autophagosome sphericities. Sphericity of one is a perfect sphere and sphericity of zero is a flat disk.

**SI Table S5. Statistics for Fig 3.**

**SI Text S6. Theoretical predictions of damped and undamped nucleation-limited growth.**

## Discussion

In this work, we explored the mechanism and dynamics of Bcl10 filament degradation, a process that is a key regulatory element in the T cell receptor-to-NF-κB signal transduction pathway. Through computational analysis of super-resolution images, we demonstrated that Bcl10 filaments are shorter in length at 40 minutes post-TCR engagement than at 20 minutes, with an increase in punctate Bcl10 and a decrease in filaments longer than 1 μm. We demonstrated that autophagosomes were preferentially in contact with Bcl10 filaments, that these contacts increased over time, that at both 20 minutes and 40 minutes post-TCR engagement there was a preference for the contacts to occur near filament ends, and that by 40 minutes post-TCR engagement, autophagosomes colocalize with Bcl10 puncta in non-random, significant manner. Together, these findings, along with our previous biochemical studies (18), imply that autophagosomes drive a degradative process that progressively shortens Bcl10 filaments, leaving behind remnants visualized as puncta which undergo slower degradation. Furthermore, we developed a stochastic Monte Carlo simulation of nucleation-limited filament growth and degradation, which we utilized to probe aspects of regulatory control of filamentous signal transduction bodies. Using this model, we ascertained that the rate of degradation was the most important element controlling the magnitude and dynamics of the response function, particularly when the homogeneity of the response has been maximized via a high activation potential and a low nucleation barrier.

Importantly, these results shed light on possible mechanisms of autophagic degradation in the poorly understood scenario in which the structures to be degraded are larger than the autophagosomes performing the degradation. We found that increased contact between autophagosomes and Bcl10 filaments corresponded to a progressive shortening of Bcl10 filament length, implying that autophagosomes degrade these large structures through piecewise degradation rather than the currently understood model of enclosure of the entire volume of a given cargo. Further, we determined that filaments to which autophagosomes attached were more likely to experience attachment near filament ends, and that degradation models that target the ends of filaments rather than the mid-region best fit the experimental observations. However, we were unable to establish whether this degradation might occur via end-based disassembly at the molecular scale, or via scission of larger regions near the end. The novel observation that the number of punctate structures in contact with autophagosomes increases significantly at 40 min vs 20 min could not be recapitulated in our model, indicating that the final degradation of Bcl10 filaments once they reach lengths <150nm and thus appear as punctae is inhibited or occurs by an independent mechanism.

The analysis methods which we developed for this study have broad applicability to computational image analysis and the study of dynamic signaling processes. Typically, radial-distribution functions are used to measure and statistically assess contacts and/or colocalizations between structures. However, because contacts between Bcl10 filaments and autophagosomes involve close-range contact and significant volumetric overlap, such an analysis was not suitable. Our image-based bootstrap-like method of random rearrangement and reanalysis of segmented objects has, to our knowledge, not been previously published, and could prove to be widely applicable in assessing colocalizations between structures in an image. Furthermore, filamentous signaling structures, particularly those whose core constituent protein contains a death domain superfamily element (as Bcl10 does), are a common motif in immunological signal transduction pathways (37–39), and many are downregulated via macroautophagy (40–42). Thus, our model of nucleation-limited filament growth and degradation, with its multiple tunable parameters that can be used to fit the model to observed imaging data, may be applicable to a variety of autophagy-regulated signal transduction pathways and reveal common mechanistic principles of immune regulation.

The simulations we developed offer intriguing insight into the regulatory control of T cell activation states. Higher values of p_activate_ in the model led to increased heterogeneity in filament lengths; since our experimental observations include heterogeneous filament lengths, it is implied that the probability of activation of Carma1 in response to TCR engagement is likewise high in biological reality. Furthermore, these higher values of p_activate_ in the model led to increased responsiveness to variations in degradation speed, along with decreased variability of the response from a variety of initial states, both outcomes that are beneficial to the overall control of T cell activation. Thus, it is probable that in activated T cells, Carma1 in the vicinity of the immunological synapse has a high likelihood of promoting Bcl10 filament nucleation, and further regulatory control of the activation potential of Carma1 would provide little overall benefit. In contrast, variations in the size of the nucleation barrier (represented by p_attach_) had little effect on the dynamics of the response, though a smaller nucleation barrier (higher p_attach_) led to slightly stabilized filaments over time and increased homogeneity of the response dynamic. Regulatory control of the nucleation barrier does not appear to provide substantial benefit to the response dynamic and is thus unlikely. Interestingly, based on the results of the simulations, governing the rate of filament degradation via control of autophagy has the greatest effect on the response dynamic. Indeed, autophagy has many regulatory control points and processes feedback from a wide variety of signaling pathways (43), providing myriad opportunities for crosstalk among T cell signaling inputs.

The results of our simulations, taken together, indicate that POLKADOTS filament growth and degradation exhibit the characteristics of an excitable system. Excitability is a phenomenon by which a system can undergo dramatic changes in behavior in response to small perturbations; components of such a system typically include non-linear thresholded/digital activation that initiates a fast, amplifying, positive feedback loop along with a slower, delayed, response-limiting negative feedback loop (44). Excitable systems characteristics are observed in a variety of biological processes, including in the polymerization and depolymerization of cytoskeletal proteins during migration (45), and in ion channels during neuronal firing cascades (46). Further, excitability is often observed when signaling networks reach a critical, near-phase-transition state (47,48), which induces dramatic changes to system dynamics, including the oligomerization of protein complexes via phase separation (49). We have previously reported that the Bcl10-dependent TCR-to-NF-κB signaling pathway is characterized by digital activation, with increased stimulus beyond a certain threshold having little effect on the signaling response of individual T cells (34). Thus, the predictions of behavior of POLKADOTS filament growth and degradation resulting from these simulations are consistent with our previous observations of the biological behavior of these signaling elements in living T cells.

Our analysis of Bcl10 filament formation and degradation suggest that this self-limiting all-or-nothing response is enabled by nucleation-limited filament assembly with a high probability of Carma1 activation in response to even minimal TCR engagement and a low nucleation barrier. The system is self-limiting via autophagy operating on filament ends, which extends the lifetime of filaments and thus signal activation (a result not inconsistent with autophagy limiting the intensity of the signal by limiting the amount of Bcl10 in POLKADOTS filaments (18)). Disassembly via autophagy results in a long refractory period due to protein degradation. More broadly, our results, when considered in combination with existing structural data (35), suggest that spatial organization of signaling components, specifically filament assembly and degradation, may be widely used in biological signaling networks to enable robust all-or-nothing (i.e., digital) responses and/or self-limiting dynamics.

## Methods

### T cell stimulation and imaging

D10.G4.1 T cells were transduced with retroviruses expressing Bcl10-GFP and TagRFP-T-LC3 (referred to as RFP-LC3) as previously described (50). Following antibiotic selection, cells were subcloned to ensure uniform imaging characteristics. The resultant D10 cell line was activated via plating on coverslips coated with anti-CD3 antibody (clone 145-2C11, BioXCell). Cells were fixed with 3% paraformaldehyde after the indicated incubation times and mounted for imaging on a custom-built instant structured illumination microscope (iSIM) utilizing a 1.45 NA oil immersion objective (36). Images were collected as z-stacks inclusive of the entire volume of the cells with a 200 nm distance between z planes. Following collection, images were deconvolved using the Richardson-Lucy algorithm as previously described (36) and converted to a new isotropically-distributed 3D coordinate grid via 3D spline interpolation.

### Segmentation of Bcl10 and LC3

Fluorescent iSIM images of Bcl10 and LC3 were binarized using an intensity threshold on a cell-by-cell and channel-by-channel basis due to fluctuations in expression and intensity. Each binarized image was convolved with a 3×3×3 kernel with values 1/3^3^ and again binarized with a threshold of (2/3)^3^ so that pixels with at least 8 occupied nearest neighbor locations were considered. All small binary pixel-noise was removed. Colocalizations between Bcl10 and LC3 were determined by identifying the voxels which directly overlap in the two binary channels.

### Skeletonization and Skeleton Measurements of Bcl10

Each of the structures in the binary Bcl10 images were individually skeletonized using a MATLAB implementation of medial-axis thinning skeletonization (51–53). Bcl10 skeletons sometimes contained erroneous spurs due to fluctuations in the volume from sensitivity to the intensity threshold and thus, filament skeletons were processed by iteratively removing spurs less than 6 voxels (150 nm) in end-to-end length, the approximate optical resolution of the iSIM. A representative example is demonstrated in **SI Fig S1**. Structurally isolated skeletons less than 6 pixels (150 nm) in length were labeled as punctate as indicated in **Fig 1**.

Where there was overlapping of multiple Bcl10 filaments, skeletons developed high-degree nodes where multiple filaments branches converged or intersected at a single branch point. This understanding of Bcl10 filament structure is inconsistent with current perspectives (14) and was treated as an imaging artifact. To estimate the appropriate skeleton segmentation in these regions, we instituted a minimum-bending-energy scheme under the assumption that filaments are less likely to develop large curvature, and thus chose the configuration that resulted in the smallest net bending energy. Bending energy for a simple elastic beam is proportional to 1/R^2^ where R is the radius of curvature of the bent region. Thus, we fit an 11-pixel segment centered at the high-degree junction between multiple skeletons to the surface of a sphere to calculate R for the segment.

Measurements from the ends of skeletons were calculated using the Floyd-Warshall algorithm where the reference point was the nearest degree-one node. Colocalizations between Bcl10 and LC3 were assigned the end-distance value from the nearest skeleton points in all overlapping voxels.

### Random Rearrangement and Resampling of Segmented Structures for Statistical Assessment

Bootstrapping is a statistical resampling procedure that allows for the estimate of a sampling distribution. Since resampling of a single fixed T cell at 20 min or 40 min post-activation is impossible, we developed an image-based resampling method to assess the sample distributions for the number and location of Bcl10-autophagosome contacts if the underlying biological processes are driven by randomness.

To carry out the procedure in this work, we identified three relevant cell structures from the 3-D super-resolution iSIM image: the cell boundary, binary Bcl10, and binary autophagosomes. For the resampling, we kept the location of the cell boundary and Bcl10 fixed. Then, we removed all autophagosomes from the image and separately saved each individual 3-D structure separately. Each 3-D autophagosome was then randomly rotated by three randomly generated angles between 0 degrees and 360 degrees about the x-, y-, and z-axis. Finally, each randomly rotated autophagosome was placed one at a time in a random location within the cell boundary. If any voxel in the currently selected autophagosome fell outside of the boundary or overlapped with an existing autophagosome, then a new location was chosen. The procedure was repeated until all autophagosomes were placed. Upon completion, the contact calculations were repeated, including the number of Bcl10-autophagosome contacts per cell, and the location of each of the contacts along connected Bcl10-filament skeletons.

### Simulations of Bcl10 Filament Growth and Degradation

Simulated cells were initialized with a variable amount of monomeric Bcl10, monomeric Carma1, and autophagosomes. To determine initial levels, each cell was assigned four random radii, R_1_, R_2_, R_3_ and R_4_, from a normal distribution with parameters μ = 1 and σ = 0.2. The initial amount of Carma1 was a_0_×R_1_^3^, Bcl10 was b_0_×R_2_^3^, autophagosomes were c_0_×R_3_^3^, and the size of the immunological synapse was d_0_×R_4_^2^, where a_0_ = 10, b_0_ = 500, c_0_ = 5, and d_0_ = 0.2, were each chosen empirically.

The simulated evolution of filament growth and degradation was controlled by the activation of Carma1 by the engaged TCR, the nucleation of Bcl10 filaments by an active Carma1, the attachment of monomeric Bcl10 to nucleated Bcl10 filaments, the attachment of autophagosomes to Bcl10 filaments, and the degradation of Bcl10 filaments by autophagosomes. Each step in the evolution was mediated by both fixed and variable probabilities.

1. Activation of Carma1 by the engaged TCR.
  a. Probability that Carma1 will collide with the TCR, p_collide_ = 0.1.
  b. Probability that the TCR was part of the immunological synapse, p_synapse_ = d_0_×R_4_^2^.
  c. Probability of Carma1 activation, p_activate_ (variable).
2. Growth of Bcl10 on activated Carma1.
  a. Probability that Bcl10 will collide with Carma1, p_collide_ = 0.01.
  b. Probability of attachment before the nucleation barrier has been reached, p_attach_ (variable).
  c. Size of the nucleation barrier (i.e., the number of attachments before the attachment probability switches from p_attach_ to p_grow_), n_barrier_ = 20.
  d. Probability of attachment after the nucleation barrier has been reached, p_grow_ = 0.4.
3. The degradation of Bcl10 by an attached autophagosome.
  a. Probability of autophagosomes colliding with a Bcl10 filament, p_collide_ = 0.1.
  b. Probability of autophagosomes attaching to a Bcl10 filament, p_attach_ = 0.1.
  c. Probability of autophagosomes degrading attached-to filaments, p_degrade_ (variable).
  d. If attached autophagosomes do not degrade, the probability that they fall off the attached-to filament, p_detach_ = 0.1.

We used physical properties of Carma1 and Bcl10 to inform the interaction probabilities in the simulations. Bcl10 is 33 kDa and its cytosolic diffusion constant is on the order of D_Bcl10_ ≈ 10 μm^2^/s, whereas Carma1 is 130 kDa and thus has a cytosolic diffusion constant on the order of D_Carma1_ ≈ 1 μm^2^/s (39). Via diffusion, the time it takes to traverse some distance d is t = d^2^/6D. T cell diameters are on the order of d_T cell_ ≈ 10 μm, and thus the time it takes for Bcl10 and Carma1 molecules to traverse the cytosol are approximately 0.2 sec and 2 sec, respectively. Thus, the collision probability for Bcl10 was chosen to be 10× larger than that of Carma1. The other rate constants, such as p_grow_ and p_attach_, have not been determined experimentally, to our knowledge.

Simulations from **Fig 4** were performed without autophagosomes and terminated upon monomeric Bcl10 being completely converted to filamentous Bcl10. Simulations from **Fig 5** were performed for either 10,000 iterations or when the number of monomeric and filamentous Bcl10 reached zero, whichever came first.

## Acknowledgements

All authors acknowledge help from Andrew York, who built the iSIM used in this study. LC and WL acknowledge valuable discussions with members of the Losert Lab and Professor John Fourkas.

## Author Contributions

**LC –** conceptualization, data curation, formal analysis, investigation (simulations), methodology, software, visualization, writing – original draft preparation

**MKT –** conceptualization, investigation (experiments), project administration, validation, visualization, writing – original draft preparation

**HS –** resources, writing – review & editing

**BCS –** conceptualization, funding acquisition, resources, supervision, writing – review & editing

**WL –** conceptualization, funding acquisition, methodology, resources, supervision, writing – review & editing

## References

1. Garner H, Visser KE de. Immune crosstalk in cancer progression and metastatic spread: a complex conversation. Nat Rev Immunol. 2020 Feb 5;1–15.

2. Theofilopoulos AN, Kono DH, Baccala R. The multiple pathways to autoimmunity. Nat Immunol. 2017 Jul;18(7):716–24.

3. Horwitz DA, Fahmy TM, Piccirillo CA, La Cava A. Rebalancing Immune Homeostasis to Treat Autoimmune Diseases. Trends Immunol. 2019 Oct;40(10):888–908.

4. Oeckinghaus A, Ghosh S. The NF-kappaB family of transcription factors and its regulation. Cold Spring Harb Perspect Biol. 2009 Oct;1(4):a000034.

5. Gilmore TD. Introduction to NF-*κ*B: players, pathways, perspectives. Oncogene. 2006 Oct;25(51):6680–4.

6. Perkins ND. Integrating cell-signalling pathways with NF-κB and IKK function. Nat Rev Mol Cell Biol. 2007 Jan;8(1):49–62.

7. Lu HY, Bauman BM, Arjunaraja S, Dorjbal B, Milner JD, Snow AL, et al. The CBM-opathies-A Rapidly Expanding Spectrum of Human Inborn Errors of Immunity Caused by Mutations in the CARD11-BCL10-MALT1 Complex. Front Immunol. 2018;9:2078.

8. Arjunaraja S, Snow AL. Gain-of-function mutations and immunodeficiency: at a loss for proper tuning of lymphocyte signaling. Curr Opin Allergy Clin Immunol. 2015 Dec;15(6):533–8.

9. Paciolla M, Pescatore A, Conte MI, Esposito E, Incoronato M, Lioi MB, et al. Rare mendelian primary immunodeficiency diseases associated with impaired NF-κB signaling. Genes Immun. 2015 Jun;16(4):239–46.

10. Herrington FD, Carmody RJ, Goodyear CS. Modulation of NF-κB Signaling as a Therapeutic Target in Autoimmunity. J Biomol Screen. 2016 Mar;21(3):223–42.

11. Sun S-C, Chang J-H, Jin J. Regulation of NF-κB in Autoimmunity. Trends Immunol. 2013 Jun;34(6):282–9.

12. Gehring T, Seeholzer T, Krappmann D. BCL10 – Bridging CARDs to Immune Activation. Front Immunol [Internet]. 2018 [cited 2020 Jan 13];9. Available from: https://www.frontiersin.org/articles/10.3389/fimmu.2018.01539/full

13. Paul S, Schaefer BC. A new look at T cell receptor signaling to nuclear factor-κB. Trends Immunol. 2013 Jun 1;34(6):269–81.

14. David L, Li Y, Ma J, Garner E, Zhang X, Wu H. Assembly mechanism of the CARMA1–BCL10– MALT1–TRAF6 signalosome. Proc Natl Acad Sci. 2018 Jan 30;201721967.

15. Paul S, Traver MK, Kashyap AK, Washington MA, Latoche JR, Schaefer BC. T Cell Receptor Signals to NF-κB Are Transmitted by a Cytosolic p62-Bcl10-Malt1-IKK Signalosome. Sci Signal. 2014 May 13;7(325):ra45–ra45.

16. Thome M, Charton JE, Pelzer C, Hailfinger S. Antigen Receptor Signaling to NF-κB via CARMA1, BCL10, and MALT1. Cold Spring Harb Perspect Biol. 2010 Sep 1;2(9):a003004.

17. Ruland J, Hartjes L. CARD–BCL-10–MALT1 signalling in protective and pathological immunity. Nat Rev Immunol. 2019 Feb;19(2):118–34.

18. Paul S, Kashyap AK, Jia W, He Y-W, Schaefer BC. Selective Autophagy of the Adaptor Protein Bcl10 Modulates T Cell Receptor Activation of NF-κB. Immunity. 2012 Jun 29;36(6):947–58.

19. Rossman JS, Stoicheva NG, Langel FD, Patterson GH, Lippincott-Schwartz J, Schaefer BC. POLKADOTS Are Foci of Functional Interactions in T-Cell Receptor–mediated Signaling to NF-κB. Mol Biol Cell. 2006 May;17(5):2166–76.

20. Qiao Q, Yang C, Zheng C, Fontán L, David L, Yu X, et al. Structural Architecture of the CARMA1/Bcl10/MALT1 Signalosome: Nucleation-Induced Filamentous Assembly. Mol Cell. 2013 Sep 26;51(6):766–79.

21. Schlauderer F, Seeholzer T, Desfosses A, Gehring T, Strauss M, Hopfner K-P, et al. Molecular architecture and regulation of BCL10-MALT1 filaments. Nat Commun. 2018 Oct 2;9(1):4041.

22. Lobry C, Lopez T, Israël A, Weil R. Negative feedback loop in T cell activation through IκB kinase-induced phosphorylation and degradation of Bcl10. Proc Natl Acad Sci. 2007 Jan 16;104(3):908–13.

23. Hu S, Du M-Q, Park S-M, Alcivar A, Qu L, Gupta S, et al. cIAP2 is a ubiquitin protein ligase for BCL10 and is dysregulated in mucosa-associated lymphoid tissue lymphomas. J Clin Invest. 2006 Jan;116(1):174–81.

24. Scharschmidt E, Wegener E, Heissmeyer V, Rao A, Krappmann D. Degradation of Bcl10 induced by T-cell activation negatively regulates NF-kappa B signaling. Mol Cell Biol. 2004 May;24(9):3860–73.

25. Wu C-J, Ashwell JD. NEMO recognition of ubiquitinated Bcl10 is required for T cell receptor-mediated NF-κB activation. Proc Natl Acad Sci. 2008 Feb 26;105(8):3023–8.

26. Zeng H, Di L, Fu G, Chen Y, Gao X, Xu L, et al. Phosphorylation of Bcl10 Negatively Regulates T-Cell Receptor-Mediated NF-κB Activation. Mol Cell Biol. 2007 Jul;27(14):5235–45.

27. Allison KA, Sajti E, Collier JG, Gosselin D, Troutman TD, Stone EL, et al. Affinity and dose of TCR engagement yield proportional enhancer and gene activity in CD4+ T cells. eLife. 2016 04;5.

28. Altan-Bonnet G, Germain RN. Modeling T cell antigen discrimination based on feedback control of digital ERK responses. PLoS Biol. 2005 Nov;3(11):e356.

29. Stefanová I, Hemmer B, Vergelli M, Martin R, Biddison WE, Germain RN. TCR ligand discrimination is enforced by competing ERK positive and SHP-1 negative feedback pathways. Nat Immunol. 2003 Mar;4(3):248–54.

30. Das J, Ho M, Zikherman J, Govern C, Yang M, Weiss A, et al. Digital Signaling and Hysteresis Characterize Ras Activation in Lymphoid Cells. Cell. 2009 Jan 23;136(2):337–51.

31. Navarro MN, Feijoo-Carnero C, Arandilla AG, Trost M, Cantrell DA. Protein kinase D2 is a digital amplifier of T cell receptor-stimulated diacylglycerol signaling in naïve CD8^+^ T cells. Sci Signal. 2014 Oct 21;7(348):ra99.

32. Huang J, Brameshuber M, Zeng X, Xie J, Li Q, Chien Y, et al. A single peptide-major histocompatibility complex ligand triggers digital cytokine secretion in CD4(+) T cells. Immunity. 2013 Nov 14;39(5):846–57.

33. Richard AC, Lun ATL, Lau WWY, Göttgens B, Marioni JC, Griffiths GM. T cell cytolytic capacity is independent of initial stimulation strength. Nat Immunol. 2018;19(8):849–58.

34. Kingeter LM, Paul S, Maynard SK, Cartwright NG, Schaefer BC. Cutting edge: TCR ligation triggers digital activation of NF-kappaB. J Immunol Baltim Md 1950. 2010 Oct 15;185(8):4520–4.

35. Qiao Q, Wu H. Supramolecular organizing centers (SMOCs) as signaling machines in innate immune activation. Sci China Life Sci. 2015 Nov;58(11):1067–72.

36. York AG, Chandris P, Nogare DD, Head J, Wawrzusin P, Fischer RS, et al. Instant super-resolution imaging in live cells and embryos via analog image processing. Nat Methods. 2013 Nov;10(11):1122–6.

37. Xu H, He X, Zheng H, Huang LJ, Hou F, Yu Z, et al. Structural basis for the prion-like MAVS filaments in antiviral innate immunity. eLife. 2014 Jan 1;3:e01489.

38. Lu A, Magupalli VG, Ruan J, Yin Q, Atianand MK, Vos MR, et al. Unified polymerization mechanism for the assembly of ASC-dependent inflammasomes. Cell. 2014 Mar 13;156(6):1193–206.

39. Moncrieffe MC, Bollschweiler D, Li B, Penczek PA, Hopkins L, Bryant CE, et al. MyD88 Death-Domain Oligomerization Determines Myddosome Architecture: Implications for Toll-like Receptor Signaling. Struct Lond Engl 1993. 2020 Mar 3;28(3):281–289.e3.

40. Tal MC, Sasai M, Lee HK, Yordy B, Shadel GS, Iwasaki A. Absence of autophagy results in reactive oxygen species-dependent amplification of RLR signaling. Proc Natl Acad Sci U S A. 2009 Feb 24;106(8):2770–5.

41. Shi C-S, Shenderov K, Huang N-N, Kabat J, Abu-Asab M, Fitzgerald KA, et al. Activation of autophagy by inflammatory signals limits IL-1β production by targeting ubiquitinated inflammasomes for destruction. Nat Immunol. 2012 Jan 29;13(3):255–63.

42. Into T, Horie T, Inomata M, Gohda J, Inoue J-I, Murakami Y, et al. Basal autophagy prevents autoactivation or enhancement of inflammatory signals by targeting monomeric MyD88. Sci Rep. 2017 21;7(1):1009.

43. Yin Z, Pascual C, Klionsky DJ. Autophagy: machinery and regulation. Microb Cell Graz Austria. 2016 Dec 1;3(12):588–96.

44. Devreotes PN, Bhattacharya S, Edwards M, Iglesias PA, Lampert T, Miao Y. Excitable Signal Transduction Networks in Directed Cell Migration. Annu Rev Cell Dev Biol. 2017;33(1):103–25.

45. Miao Y, Bhattacharya S, Edwards M, Cai H, Inoue T, Iglesias PA, et al. Altering the threshold of an excitable signal transduction network changes cell migratory modes. Nat Cell Biol. 2017 Apr;19(4):329–40.

46. Goldbeter A, Dupont G, Berridge MJ. Minimal model for signal-induced Ca2+ oscillations and for their frequency encoding through protein phosphorylation. Proc Natl Acad Sci. 1990 Feb 1;87(4):1461–5.

47. Kinouchi O, Copelli M. Optimal dynamical range of excitable networks at criticality. Nat Phys. 2006 May;2(5):348–51.

48. Ma Z, Turrigiano GG, Wessel R, Hengen KB. Cortical Circuit Dynamics Are Homeostatically Tuned to Criticality In Vivo. Neuron. 2019 Nov 20;104(4):655–664.e4.

49. Dine E, Gil AA, Uribe G, Brangwynne CP, Toettcher JE. Protein Phase Separation Provides Long-Term Memory of Transient Spatial Stimuli. Cell Syst. 2018 Jun 27;6(6):655–663.e5.

50. Traver MK, Paul S, Schaefer BC. T Cell Receptor Activation of NF-κB in Effector T Cells: Visualizing Signaling Events Within and Beyond the Cytoplasmic Domain of the Immunological Synapse. Methods Mol Biol Clifton NJ. 2017;1584:101–27.

51. Blum H. A Transform for Extracting New Descriptors of Shape. 1967;362–80.

52. Lee TC, Kashyap RL, Chu CN. Building Skeleton Models via 3-D Medial Surface Axis Thinning Algorithms. CVGIP Graph Models Image Process. 1994 Nov 1;56(6):462–78.

53. Kollmannsberger P, Kerschnitzki M, Repp F, Wagermaier W, Weinkamer R, Fratzl P. The small world of osteocytes: connectomics of the lacuno-canalicular network in bone. New J Phys. 2017 Jul;19(7):073019.

